# Transcriptome profiling of histone writers/erasers enzymes across spermatogenesis, mature sperm and pre-cleavage embryo: Implications in paternal epigenome transitions and inheritance mechanisms

**DOI:** 10.1101/2022.05.29.493915

**Authors:** Gastón Barbero, Maximiliano de Sousa Serro, Camila Perez Lujan, Alfredo D. Vitullo, Candela R. González, Betina González

**Author notes:** These authors share last authorship (Candela R. González, Betina González).

## Abstract

Accumulating evidence points out that sperm carry epigenetic instructions to embryo in the form of retained histones marks and RNA cargo that can transmit metabolic and behavioral traits to offspring. However, the mechanisms behind epigenetic inheritance of paternal environment are still poorly understood. Here, we curated male germ cells RNA-seq data and analyzed the expression profile of all known histone lysine writers and erasers enzymes across spermatogenesis, unraveling the developmental windows at which they are upregulated, and the specific activity related to canonical and non-canonical histone marks deposition and removal. We also characterized the epigenetic enzymes signature in the mature sperm RNA cargo, showing most of them positive translation at pre-cleavage zygote, suggesting that paternally-derived enzymes mRNA cooperate with maternal factors to embryo chromatin assembly. Our study shows several histone modifying enzymes not described yet in spermatogenesis and even more, important mechanistic aspects behind transgenerational epigenetics. Epigenetic enzymes not only can respond to environmental stressors, but could function as vectors of epigenetic information and participate in chromatin organization during maternal-to-zygote transition.

## INTRODUCTION

In the last years, accumulating research has documented that male germ cell maturation presents windows of vulnerability for epigenetic reprogramming by environmental stressors that can affect fertility and even transmit developmental, metabolic and behavioral traits to offspring (Bale, 2015). This type of non-genetic “Lamarckian” transmission has been established for paternal lifestyle and different forms of chronic stress, drug abuse, dietary change, and social defeat, involving changes in non-coding RNAs cargo, DNA methylation, and histone post-translational modifications (PTMs) (Bale, 2105). However, most of the research in epigenetic inheritance has documented the role of non-coding RNAs and DNA methylation patterns, whereas the mechanisms by which the environment can trigger specific changes on histone PTMs are still largely unexplored.

Spermatogenesis is a finely regulated process where unipotent spermatogonia undergo meiosis and, subsequently, spermiogenesis, to become sperm cells. During the developmental stages of spermatogenesis, male germ cells experience dramatic chromatin reorganization, where approximately 90% (human) to 95% (mouse) of histones are evicted and replaced by protamines to compact the paternal genome (Rajender et al., 2011). Importantly, a small fraction of histones in the sperm genome is retained in specific locations and carries several PTMs that play critical roles in epigenetic regulation of spermatogenesis and early embryonic development (Luense et al., 2016). The epigenetic program of histone PTMs that controls spermatogenesis is driven by enzymatic families known as writers and erasers of the so-called “histone code”, that modulate chromatin structure and transcription (Kouzarides, 2007). These epigenetic enzymes catalyze the deposition or removal of specific PTMs such as acetylation, methylation, phosphorylation, crotonylation, among others, at specific amino acid residues on the nucleosome’s core histones H2A, H2B, H3, and H4, and also on the linker H1/H5 (Luense et al., 2016). In male germ cells, histone PTMs not only control active vs repressed chromatin states but also a range of processes including DNA replication and repair, chromosome maintenance and histone eviction (Luense et al., 2016). So far, the most studied histone PTMs related to the control of germ cells mitosis, meiosis and spermiogenesis are methylated and acetylated lysines (K) on H2A/B, H3 and H4. Histone K acetylation can be dynamically regulated by the opposing action of acetyltransferases (HATs) and deacetylases (HDACs). Acetylation of K residues neutralizes the positive charge on histones, allowing DNA-binding proteins better access to the DNA and resulting in activation of gene expression as well as histone eviction (Rajender et al., 2011). Unlike acetylation, methylation does not affect histone charge but regulates recognition and interaction with chromatin-binding proteins that control the transcription or respond to DNA damage (Rajender et al., 2011). Histone K methylation is finely regulated by methyltransferases (KMTs) and demethylases (KDMs) that control the mono-, di-, and/or tri-methylation of specific residues, and this can either activate or repress transcription depending on the residue position, the number of methylations and the presence of other methyl or acetyl groups in the vicinity (Mosammaparast and Shi, 2010).

The role of epigenetic enzymes comes to focus, as they are responsible for the histones’ PTMs writing and erasing balance. Moreover, histone-modifying enzymes have been shown to respond to the organism physiology and metabolism, several disease conditions, and environmental stressors (Bale, 2015; Nebbioso et al., 2018). Even though epigenetic enzymes are the bridge linking the environment with epigenetic inheritance through retained histones PTMs, an in-depth analysis of their expression patterns throughout the spermatogenic process is still lacking. Moreover, these enzymes provide new targets for therapies for numerous diseases (Prachayasittikul et al., 2017) but the potential effect of these compounds on epigenetic inheritance is generally overlooked. To further elucidate the mechanisms driving the histone PTMs during spermatogenesis, a better understanding of the gene expression profiles of epigenetic modifying enzymes in male germ cells is required. Here, we curated transcriptomic data from spermatogonia to mature sperm populations and analyzed the expression profile and dynamic changes of all the known families of histone K acetylation and methylation writers and erasers. We provide a complete picture of the epigenetic enzymes across spermatogenesis in mice and the developmental windows when the transcription of these enzymes may be more susceptible to environmental disruption. Moreover, we analyzed pre-cleavage zygote translatome, confirming that mRNAs of several histone modifying enzymes carried by sperm are associated with ribosomes for protein synthesis in 1-cell embryo. Not only do we expand the knowledge on epigenetic enzymes with recognized roles in spermatogenesis, but provide evidence of many more whose roles in male germ cell development and zygote have not been described yet.

## METHODS

### Data retrieval

FASTQ files were downloaded from NCBI GEO and ArrayExpress databases. Transcriptomic data of male germ cells was obtained from GSE162740, consisting of germ cells isolated from “Stra8-Tom” mice obtained from the cross of Stra8-cre mice with CAG-LSLtdTomato mice (Mayorek et al., 2022). The gradient expression of the Tomato transgene under Stra8 expression profile, combined with immunostaining for cKit to further discriminate the SGund and SGdiff spermatogonia, was used to FACS sort 6 populations of germ cells in triplicates, and sequence in an Illumina HiSeq 2500 platform. Sperm cells data were obtained from: i) GSE81216, that sampled 2 total sperm and 2 sperm heads from C57BL/6J (JAX) mice using Ion Torrent PGM platform, ii) GSE88732, that sampled 4 mature sperm from adult C57BL/6J mice using Illumina HiSeq 4000 platform, and iii) E-MTAB-5834, that sampled 4 control mature sperm from adult C57BL/6J mice using Illumina HiSeq 2500 platform. Data from one cell embryo, 2 total and 2 ribosome-bound RNA RPKMs, were obtained from GSE169632.

### Data processing

FASTQ files from germ cells and sperm were processed with the following pipeline: quality control with FASTQC, trimming with fastp, mapping with STAR and GRCm39 (MM10 gencode.vM29.annotation), and counting with featureCounts. Processed data was manipulated using Rstudio (v1.2.1033) and Tidyverse packages Dplyr and ggplot2 (Wickham et al., 2019). The male germ cells count matrix was analyzed with DESeq2 package (Love et al., 2014), using the likelihood ratio test (LRT) for longitudinal data instead of the Wald test, and a formula that accounted for batch and cell group effects. Comparisons between consecutive cells groups were obtained by contrasts. Counts were converted to RPKM using mean gene length extracted from the gencode.vM29 file. Hierarchical clustering was performed with pheatmap package. The sperm RPKMs obtained for the three selected datasets were converted to percentile rank, and the sperm mean percentile rank calculated. Word cloud plot was performed with ggwordcloud package. The embryo traslational eficiency (TE) was calculated as Ribo-seq RPKM / total RPKM for each gene, with RPKM > 0.5. All the plots shown in this study can be reproduced by downloading the counts tables, enzymes metadata table and R scripts available at https://github.com/Gonzalez-Lab/Gonzalez-2022-germ-cells.

## RESULTS

### Epigenetic enzymes and histone K acetylation and methylation marks distribution during spermatogenic stages

Table 1 shows a curated list of writers and erasers families for histone K acetylation and methylation obtained from the Uniprot database, as proteins with confirmed catalytic activity towards histones K residues, and their most representative K targets reported in mammalian cells. Open chromatin is characterized by the presence of acetylated histones at several K residues such as H2AK5/9/13/15ac, H2BK5/12/15/20/23/24ac, H3K4/9/14/18/27/36/56/79/122ac, and H4K5/8/12/16/20/79/91ac. Many of these sites are also targets of mutually exclusive methylation. Figure 1A shows the canonical histone methylation sites that are found on H3K4/9/27/36/79 and H4K20, and characteristic of active or repressed chromatin. For instance, H3K4me1/2/3, H3K36me3, H3K9me1, H3K27me1, H3K79me2/3, and H4K20me1 are found in enhancers, promoters, and gene bodies of active genes, whereas H3K9me2/3, H3K27me2/3, and H4K20me3 are found in heterochromatin, telomeric regions and inactivated X chromosome (Fig. 1A) (Black et al., 2012). In addition, there are multiple non-canonical histone methylation sites on H2AZK7, H3K23/56/63 and H4K5/12 with unknown functions, except for H3K56me3 that is involved in heterochromatin formation (Black et al., 2012). In Figure 1B we summarized the available information on the most important histone K acetylation and methylation marks distribution during spermatogenic stages, obtained from reported immunohistochemical and/or proteomic studies. We observed two main profiles on the marks distribution across spermatogenesis: H3/4ac, H3K4me2/3 and H3K9me2/3 increase with spermatogonia differentiation and early meiosis, decrease at late meiosis, and regain high levels at spermiogenesis, whereas H3K36me2/3, H3K27me2/3, H3K79me3 and H4K20me3 show sustained increased expression towards differentiation, but H4K20me is erased at RStid stage.

**Table 1.**
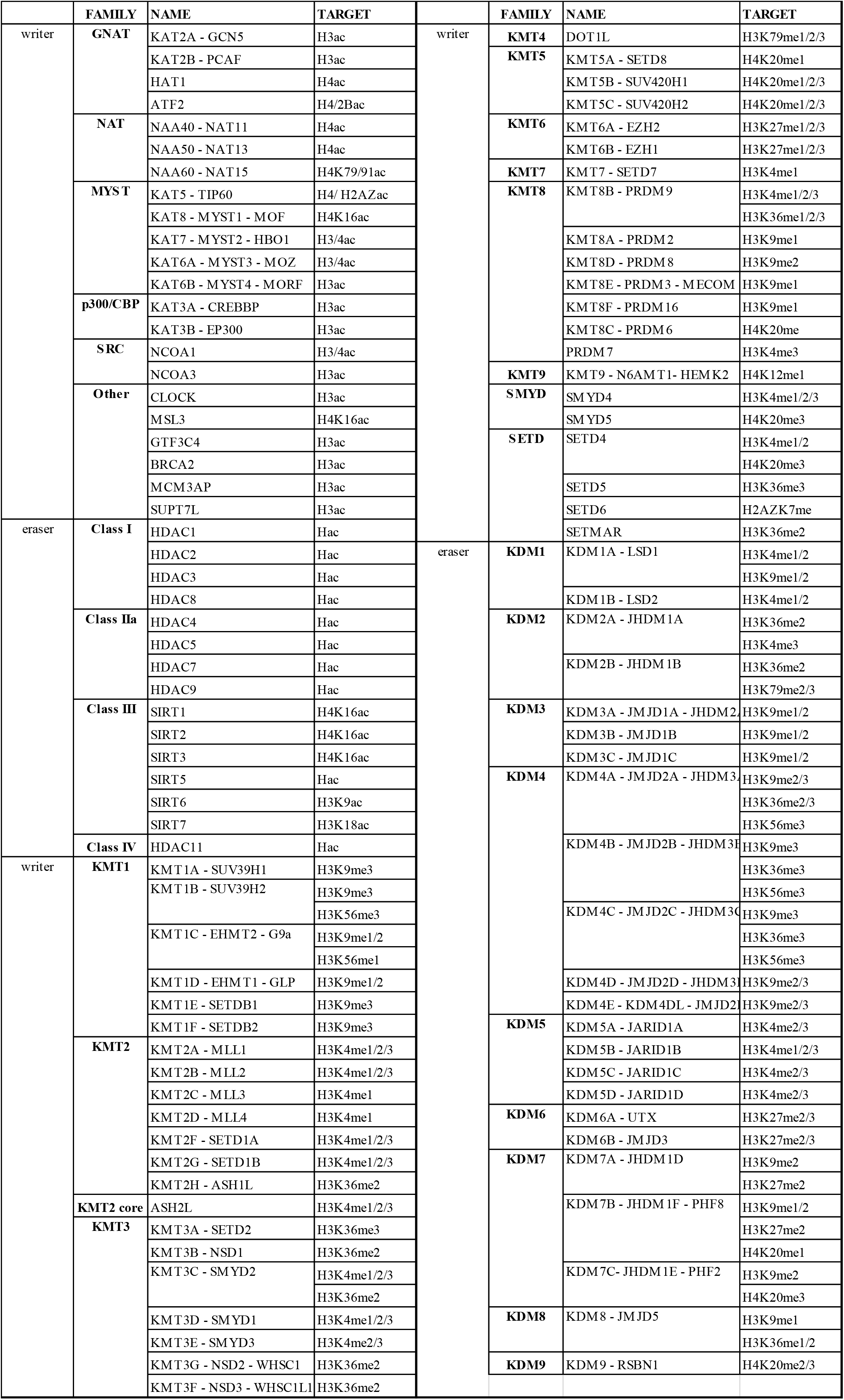
Writers and erasers families for histone K acetylation and methylation and the targets described. The HATs can be divided into six major families: 1) the GNATs (GCN5-related N-acetyltransferases), 2) the NATs (N-terminal acetylases, which also possess a GNAT domain), 3) the MYST, 4) the p300/CBP, 5) the SRC steroid receptors coactivators, and 6) other HATs (Wapenaar and Dekker, 2016). HATs actions are counteracted by the HDACs, which include zinc-dependent class I, class IIa, and class IV HDACs, and NAD-dependent class III Sirtuins (Wapenaar and Dekker, 2016). On the other hand, the KMTs present two domains with annotated lysine methyltransferase activity: 1) SET domains, that catalyze all the canonical methylation sites except for H3K79, and 2) 7βS domains (DOT1L and N6AMT1) (Mosammaparast and Shi, 2010). The KDMs can be divided according to their catalytic activity in: 1) amine oxidases (KDM1), and 2) Jumonji C demethylases (KDM2/3/4/5/6/7/8) (Mosammaparast and Shi, 2010).

**Figure 1.**
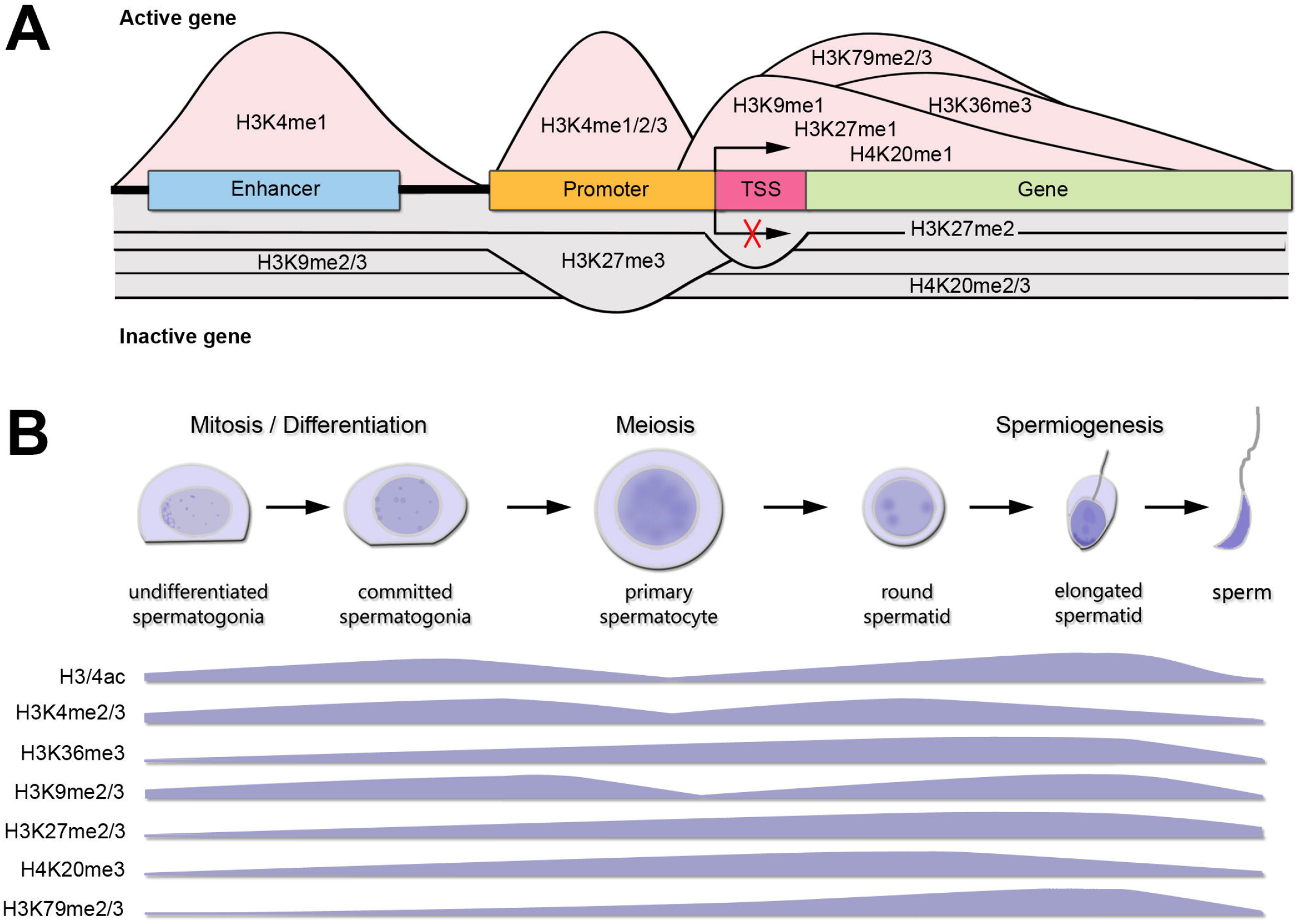
Canonical histone K marks distribution during spermatogenesis. A) Histone K methylation PTMs distribution at DNA elements and functional roles in gene expression for an active gene (pink shading) or inactive gene (gray shading). TSS: transcription start site. Only K modifications with sufficient and consensual information about distribution across genes are shown. B) Levels of main histone K PTMs involved in chromatin remodeling during spermatogenesis. The data presented is summarized from the following references: Song et al., 2011; Godmann et al., 2007; Zuo et al., 2018; Luense et al., 2016; Wang et al., 2021. SGund: spermatogonia Kit-, SGdiff: spermatogonia Kit+, PreL: pre-leptotene spermatocytes, LZ: leptotene/zygotene spermatocytes, PD: pachytene/diplotene spermatocytes, RStid: round spermatid.

### Histone K acetylation and methylation enzymes expression during spermatogenesis

To analyze the expression profiles of writers and erasers families for histone K acetylation and methylation across spermatogenesis, we curated RNA-seq data available in purified male germ cells from adult mice and selected the dataset GSE162740 (Mayorek et al., 2022). This dataset contains the mRNA profiles of 6 populations of germ cells isolated in triplicates: undifferentiated (SGund) and differentiated spermatogonia (SGdiff) (spermatogonial phase), primary spermatocytes, including pre-leptotene (PreL), leptotene/zygotene (LZ) and pachytene/diplotene (PD) stages (meiosis), and round spermatids (RStid) (spermiogenesis phase). We downloaded the FASTQ files and mapped them to MM10 genome using STAR and featureCounts to obtain the count matrix. We corroborated the cell stages and purity of the dataset by analyzing spermatogenic and somatic cell marker genes profiles in each sample (Fig. S1 and Table S1). Figure 2 shows the hierarchical clustering of z-scores for the 102 epigenetic enzymes listed in Table 1. Column clustering shows two main profiles between the spermatogonial - early meiotic vs late meiotic - spermiogenic phases. SGdiff and PreL stages are the most similar, whereas SGund and LZ show large clusters of enzymes upregulation. The late meiotic and spermiogenesis genes show two main clusters: genes specifically upregulated at PD and genes that increase at PD and remain high or even increase expression at RStid stage.

**Figure 2.**
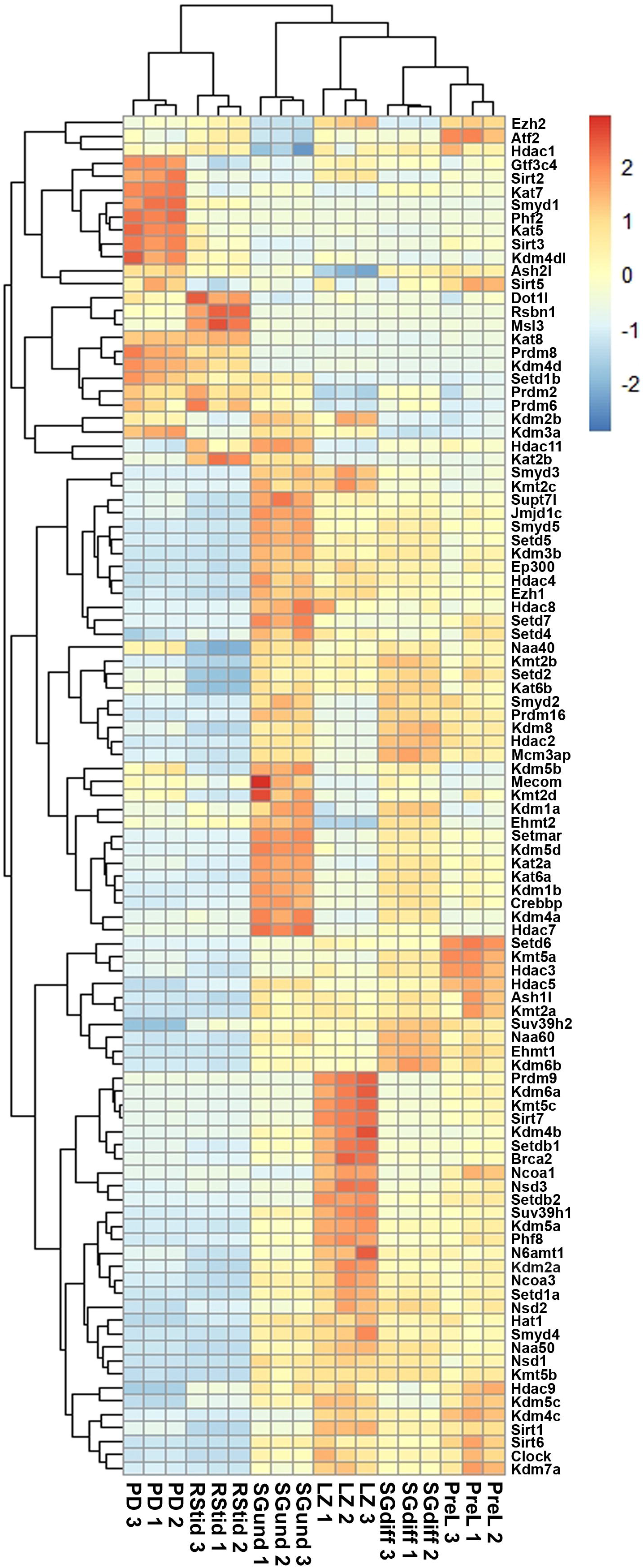
Unsupervised hierarchical clustering analysis of male germ cells transcriptome. Heatmap showing z-score values of histone K acetylation and methylation writer/eraser enzymes. SGund: spermatogonia Kit-, SGdiff: spermatogonia Kit+, PreL: pre-leptotene spermatocytes, LZ: leptotene/zygotene spermatocytes, PD: pachytene/diplotene spermatocytes, RStid: round spermatid.

### Epigenetic enzymes mRNA regulation across spermatogenesis

To analyze upregulated mRNAs of epigenetic enzymes across male germ cells differentiation, we performed differential gene expression (DEG) analysis using the DESeq2 algorithm, and performed contrasts on consecutive stages of cell development: SGdiff vs SGund, PreL vs SGdiff, LZ vs PreL, PD vs LZ and RStid vs PD (Fig. 3A). For each developmental stage, we constructed heatmaps for the upregulated enzymes (padj<0.05, log2FC>0.5) during the spermatogonial (Fig. 4), meiosis prophase I (Fig. 5) and spermiogenesis (Fig. 6) phases. Figure 3B shows the enzymes not significantly upregulated at any cell stage. These epigenetic enzymes showed a pattern of high expression at the spermatogonial phase that decreased with cell differentiation, and will be considered as characteristic of the spermatogonial - early meiotic phase. To visualize the functional profile of the upregulated enzymes at each stage, we constructed dot plots indicating the target histone mark, the writer/eraser nature and mRNA abundance (RPKM).

**Figure 3.**
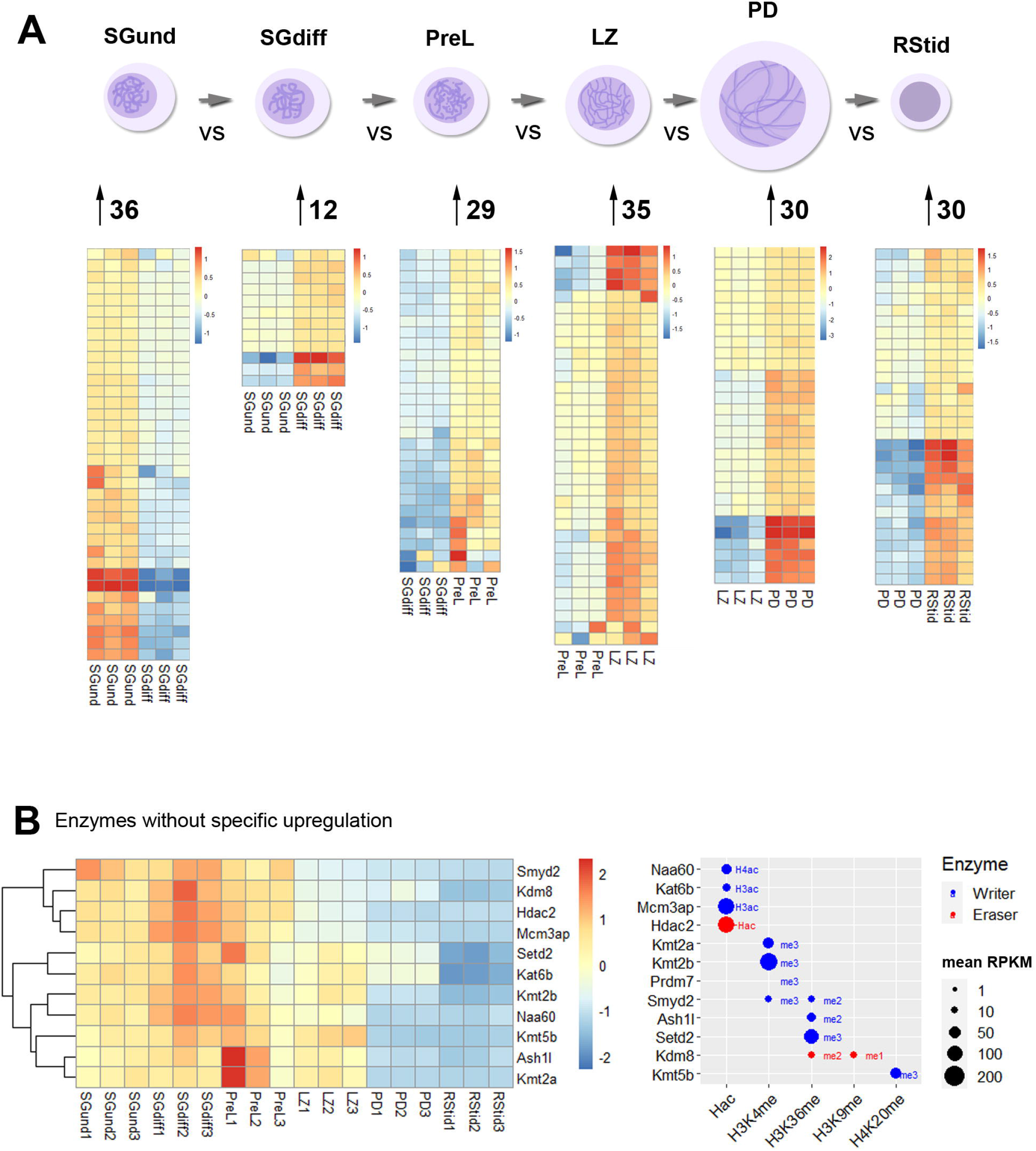
Histone K acetylation and methylation epigenetic enzymes expression during spermatogenesis. A) Number of upregulated epigenetic enzymes and heatmap showing z-score values for each germ cell stage. Differential gene expression analysis was conducted with DESeq2 with LRT test and contrasts on consecutive populations across differentiation (SGdiff vs SGund, PreL vs SGdiff, LZ vs PreL, PD vs LZ and RStid vs PD). Upregulated genes were selected with padj<0.05 and log2FC>0.5. B) Left: Heatmap showing epigenetic enzymes not significantly upregulated at any germ cell stage across spermatogenesis. Right: Dot plot showing enzymes mean RPKM values from SGund to PD populations. Blue dots: writers, red dots: erasers. SGund: spermatogonia Kit-, SGdiff: spermatogonia Kit+, PreL: pre-leptotene spermatocytes, LZ: leptotene/zygotene spermatocytes, PD:pachytenee/diplotene spermatocytes, RStid: round spermatid. The H2/3/4ac reference indicates the histone target reported for each lysine acetylation writer. Hac indicates that lysine acetylation eraser was reported to erase acetylation on all histones lysines.

**Figure 4.**
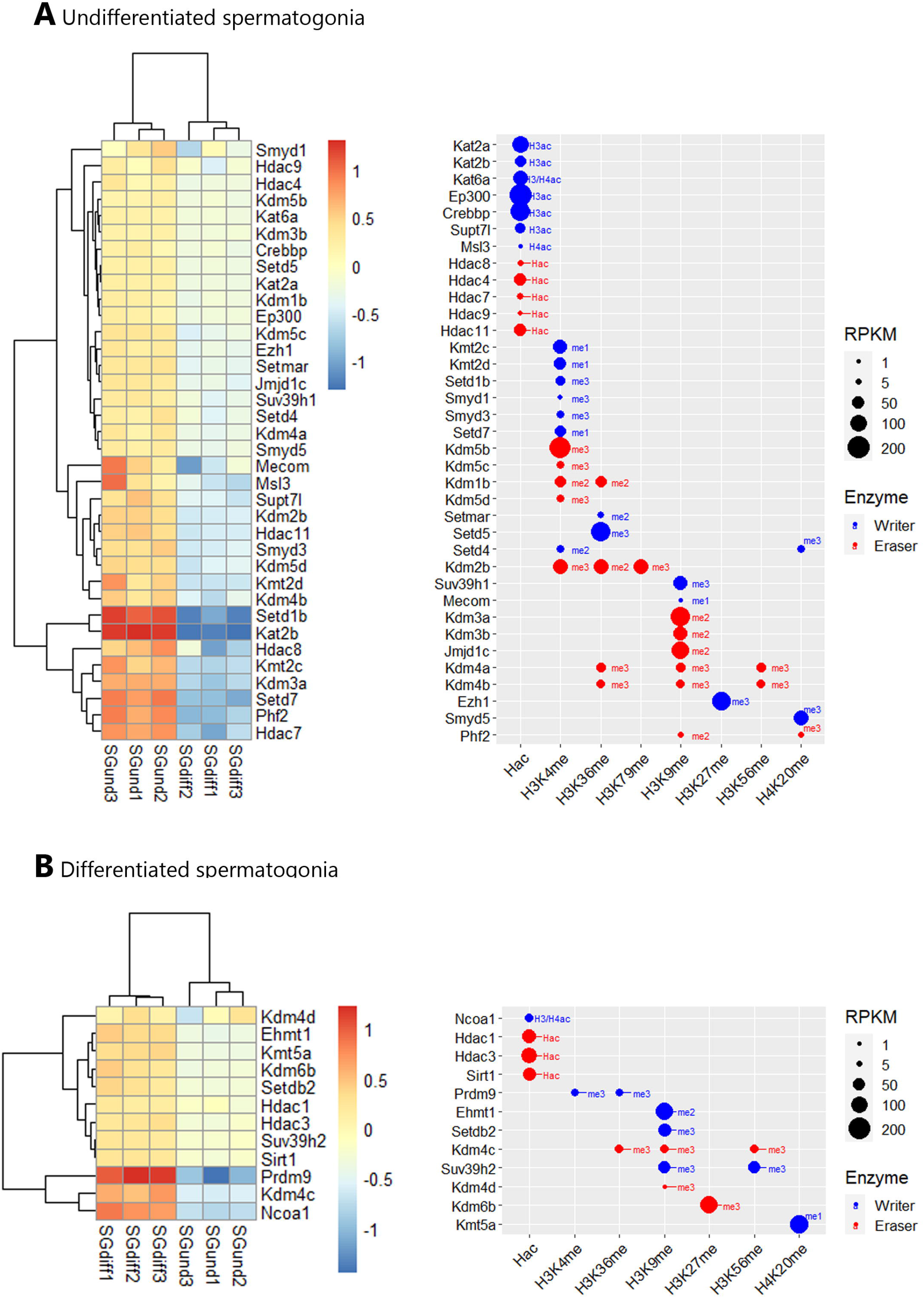
Epigenetic histone K writers and erasers upregulation at spermatogonial phase. The left panel indicates the heatmap showing z score values and the right panel indicates the Dot plot showing enzymes RPKM values in SGund (A) and SGdiff (B) population. Blue dots: writers, red dots: erasers. SGund: spermatogonia Kit-, SGdiff: spermatogonia Kit+. The me1/2/3 reference for each dot indicates the highest methyl position reported for each lysine methylation enzyme. The H2/3/4ac reference indicates the histone target reported for each lysine acetylation writer. Hac indicates that lysine acetylation eraser was reported to erase acetylation on all histones lysines.

**Figure 5.**
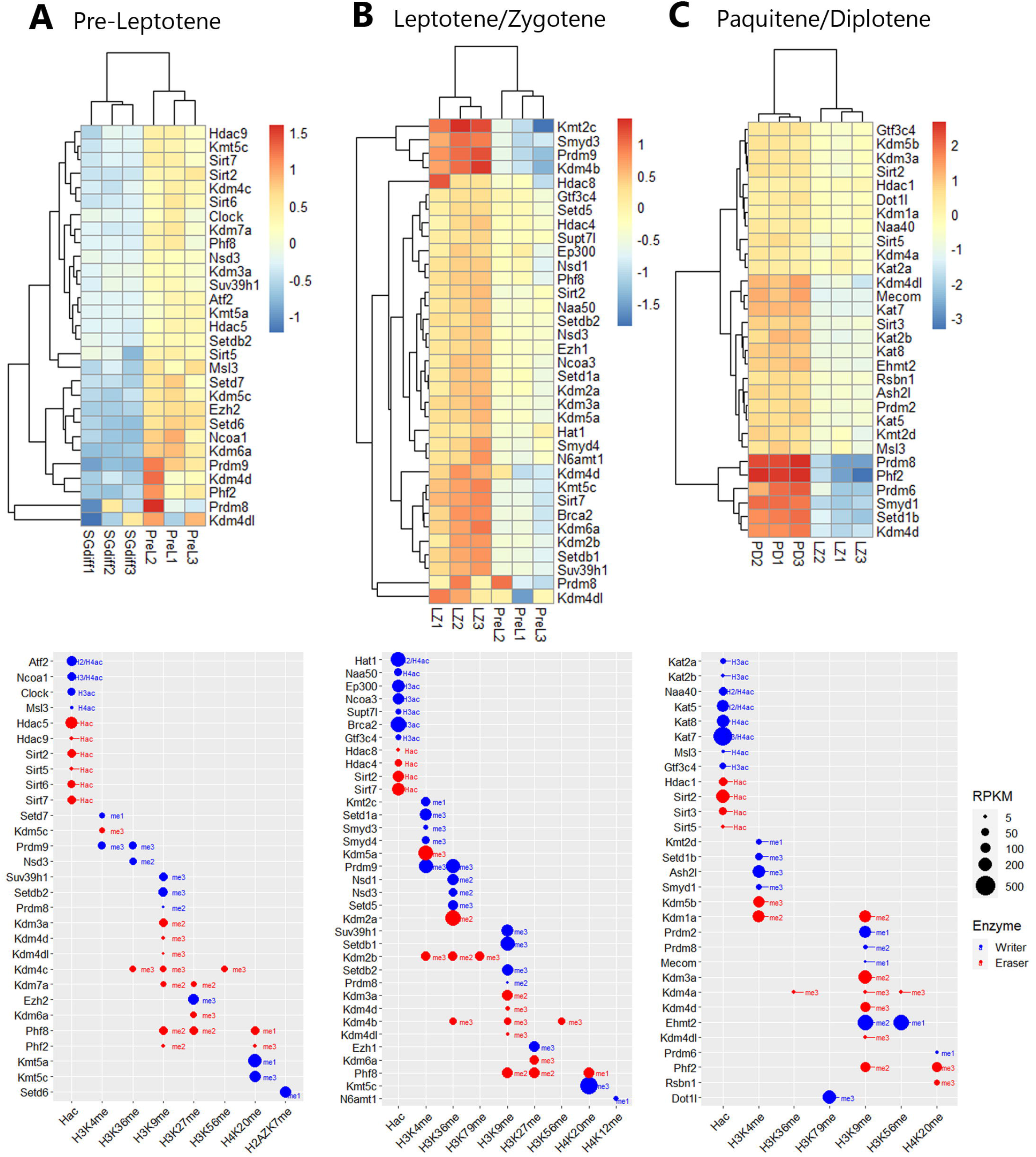
Epigenetic histone K writers and erasers upregulation at meiosis prophase I. The left panel indicates the heatmap showing z score values and the right panel indicates the Dot plot showing enzymes RPKM values, in PreL (A), LZ (B) and PD (C) spermatocytes. Blue dots: writers, red dots: erasers. PreL: pre-leptotene spermatocytes, LZ: leptotene/zygotene spermatocytes, PD:pachytenee/diplotene spermatocytes. The me1/2/3 reference for each dot indicates the highest methyl position reported for each lysine methylation enzyme. The H2/3/4ac reference indicates the histone target reported for each lysine acetylation writer. Hac indicates that lysine acetylation eraser was reported to erase acetylation on all histones lysines.

**Figure 6.**
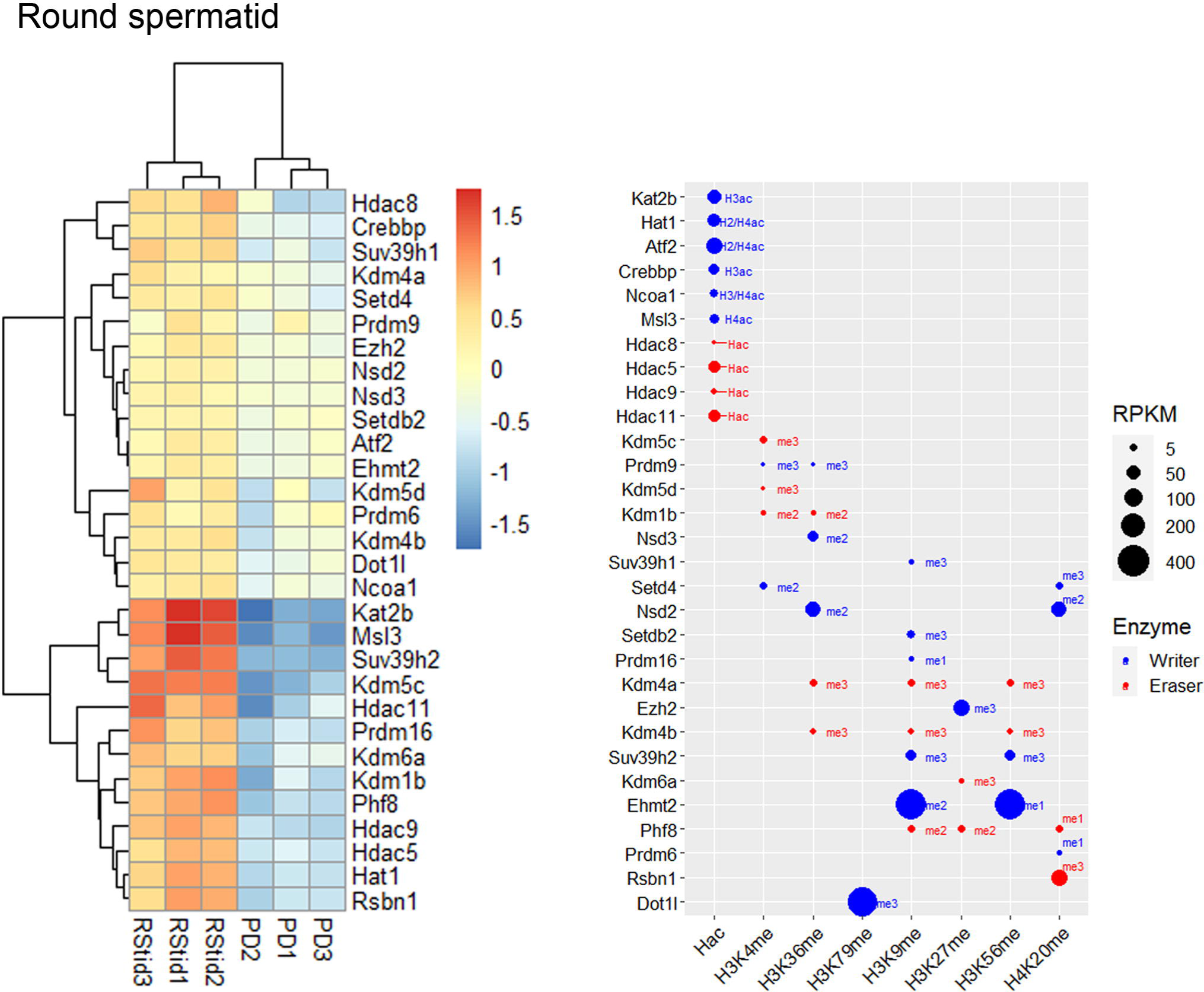
Epigenetic histone K writers and erasers upregulation at spermiogenesis. The left panel indicates the heatmap showing z score values and the right panel indicates the Dot plot showing enzymes RPKM values in RStid. Blue dots: writers, red dots: erasers. PD: pachytene/diplotene spermatocytes, RStid: round spermatid. The me1/2/3 reference for each dot indicates the highest methyl position reported for each lysine methylation enzyme. The H2/3/4ac reference indicates the histone target reported for each lysine acetylation writer. Hac indicates that lysine acetylation eraser was reported to erase acetylation on all histones lysines.

### Expression profile of histone K acetylation and methylation enzymes during spermatogenesis phases

During the spermatogonial phase, the SGund population is characterized by a mitotic activity to maintain testicular homeostasis through self-renewal. Some of these SGund initiate extensive chromatin reorganization and their epigenetic landscape shifts dramatically to differentiate to committed SGdiff (Shirakawa et al., 2013). SGund showed increased KDM3/4 H3K9me erasers, and SGdiff increased KMT1 H3K9/K56me and KMT5 H4K20me writers expression (Fig. 4). At this phase, H3K27me regulation showed high expression of the polycomb H3K27me writer subunit *Ezh1* at the SGund, and the transition to SGdiff upregulated the H3K27me eraser *Kdm6b*. We found enzymes with high expression at both SGund and SGdiff, including important methylation writers such as K*mt2a/b* (*Mll1/2*), *Setd2, Kmt5b* (*Suv420h1*), *Ehmt2* and *Kdm1a* (Fig. 2 and Fig. 3B). The SGund also showed high levels of several writers and erasers for H3K4me and H3K36me, including COMPASS subunits *Kmt2c/d* (*Mll3/4*) and *Setd1b*, the H3K36me3 writer *Setd5*, and erasers *Kdm1b* and polycomb PRC1 subunit *Kdm2b*. Several HATs showed increased levels at SGund, including *Kat2a/b, Ep300, Msl3* and *Supt7l*. The SGund also showed high expression of class I *Hdac8*, class II *Hdac4/7/9* and class IV *Hdac11*. The transition to SGdiff specifically upregulated H3K4me and H3K36me *Prdm9* and *Nsd2* writers, together with increased *Ncoa1*, class I *Hdac1/3* and class III *Sirt1*. Other HATs like *Kat6a/b, Naa60* and *Mcm3ap* were elevated at both spermatogonial populations (Fig. 3B).

Meiotic entry occurs concomitantly with the pre-meiotic S phase and is followed by the meiotic prophase that initiates with PreL stage. Here, we found some enzymes that continue to increase expression from SGdiff to LZ stage, including *Prdm9, Setdb2* and *Kmt5a/c*, and enzymes that seem to be specific to each meiotic stage, like *Setd6* at PreL stage that deposits H2AZK7me1 mark and *N6amt1* at LZ stage, that deposits H4K12me mark (Fig. 5). Histone acetylation writers increased from PreL to PD stage, with upregulated expression of almost all members of GNATs, NATs, SRC, MYST, and p300/CBP families, and other HATs like *Supt7l, Msl3, Clock, Gtf3c4* and *Brca2*, showing most of them high RPKM values. During meiotic prophase I, several class I, II and III HDACs were increased, including *Hdac1/8, Hdac4/5/9* and *Sirt2/3/5/6/7*. Additionally, note that the major enzymatic regulation of active H3K4me and H3K36me marks occurs during chromosomal synapsis at LZ stage, allowing the crossing over of homologous chromosomes at hotspots during PD stage. Moreover, H3K36me writers *Nsd1/2/3, Setd5* and *Prdm9* peak expression from PreL to LZ stage, while PD stage did not show peak expression of neither H3K36me writers nor erasers.

Also, prophase I stages showed increased expression of KDM5 family and *Kdm1a*, consistent with the decreased H3K4me2/3 global expression reported at this stage (Fig. 1B). We also detected increased expression and high RPKM values of H3K27me writers and erasers at PreL and LZ stages, including polycomb writers *Ezh1/2*, and *Kdm6a*, which form the COMPASS complex with *Kmt2b/c/d* and *Ashl2* H3K4me writers increased at these stages. Meiotic prophase I also showed upregulation of KDM7 family, *Kdm7a* and *Phf2/8*, that mediate gene activation programs by removal of several histone repressive marks. In addition, there is considerable enzymatic regulation on H3K9me mark, evidenced in the number and peak expression of H3K9me writers and erasers during all stages of prophase I, including KDM3/4 families that would mediate the decreased global H3K9me3 levels detected at this stage (Fig. 1B).

Spermiogenesis is the final phase of spermatogenesis, where the haploid RStids engage in chromatin remodeling programs involving specific histone variants incorporation, H3/4 hyperacetylation and subsequent replacement of 90-99% histones by protamines (PRMs) to compact the nucleus. Figures 6 shows that the RStid stage presents expression of histone K acetylation writers and erasers that continued to increase expression from PD stage, including *Kdm4a, Msl3, Ehmt2, Rsbn1* and *Dot1l*, and others that seemed to reactivate expression from previous spermatogenic and early meiotic stages. We found specifically upregulation of several HATs at RStids, and also *Kat8* that maintained high levels from PD stage (Fig. 2) consistent with the histone hyperacetylation that takes place at this stage (Fig. 1B). The RStid stage showed high RPKM values for the H3K79me writer *Dot1l*, the H3K27me writer *Ezh2* and the H3K9me2 writer *Ehmt2*, consistent with the high global expression of these marks at spermiogenesis stage (Fig. 1B). Additionally, RStid population showed peak expression of *Rsbn1*, a known specific eraser involved in decreasing H4K20me levels at spermiogenesis (Fig. 1B).

### Epigenetic enzymes detected at mature sperm and 1 cell embryo translatome

To characterize the epigenetic enzymes mRNAs present in the mature sperm, we analyzed data from three datasets that performed long non-coding and mRNA RNAseq: GSE81216 (total and head sperm, Schuster et al., 2016), GSE88732 (Zhang et al., 2017), and E-MTAB-5834 (Gapp et al., 2020). For each study, we obtained the mean RPKM for each gene and converted it to percent rank to transform values into a 0-1 scale, where 0 means no detection and 1 is the highest detected gene in each study (see Supplemental Table S2). Then, we calculated the mean percent rank for the three studies. Figure 7A shows a positive correlation between sperm and RStid mean percent rank, which favors the idea that the sperm enzymes mRNA comes from the spermiogenesis phase. In addition, with the aim to elucidate if the epigenetic enzymes detected in mature sperm may participate in embryo chromatin assembly, we analyzed recently reported data that profiled the mRNA translation landscape in mouse pre-implantation embryos by Ribo-seq (Zhang et al., 2022). We analyzed the translation efficiency (TE), obtained as mean RPKM detected at ribosomes/mean RPKM detected at whole RNAseq (RPKM>0.5) (see Supplemental Table S3). Figure 7B shows a word cloud plot where word size indicates enzyme abundance (mean percent rank) in mature sperm, and red color indicates positive translation detected at 1 cell embryo. Figure 7C shows that several writers and erasers of all histone acetylation and methylation marks are detected in the translatome of the 1-cell embryo, where many of these mRNAs are highly detected in the sperm RNA pool.

**Figure 7.**
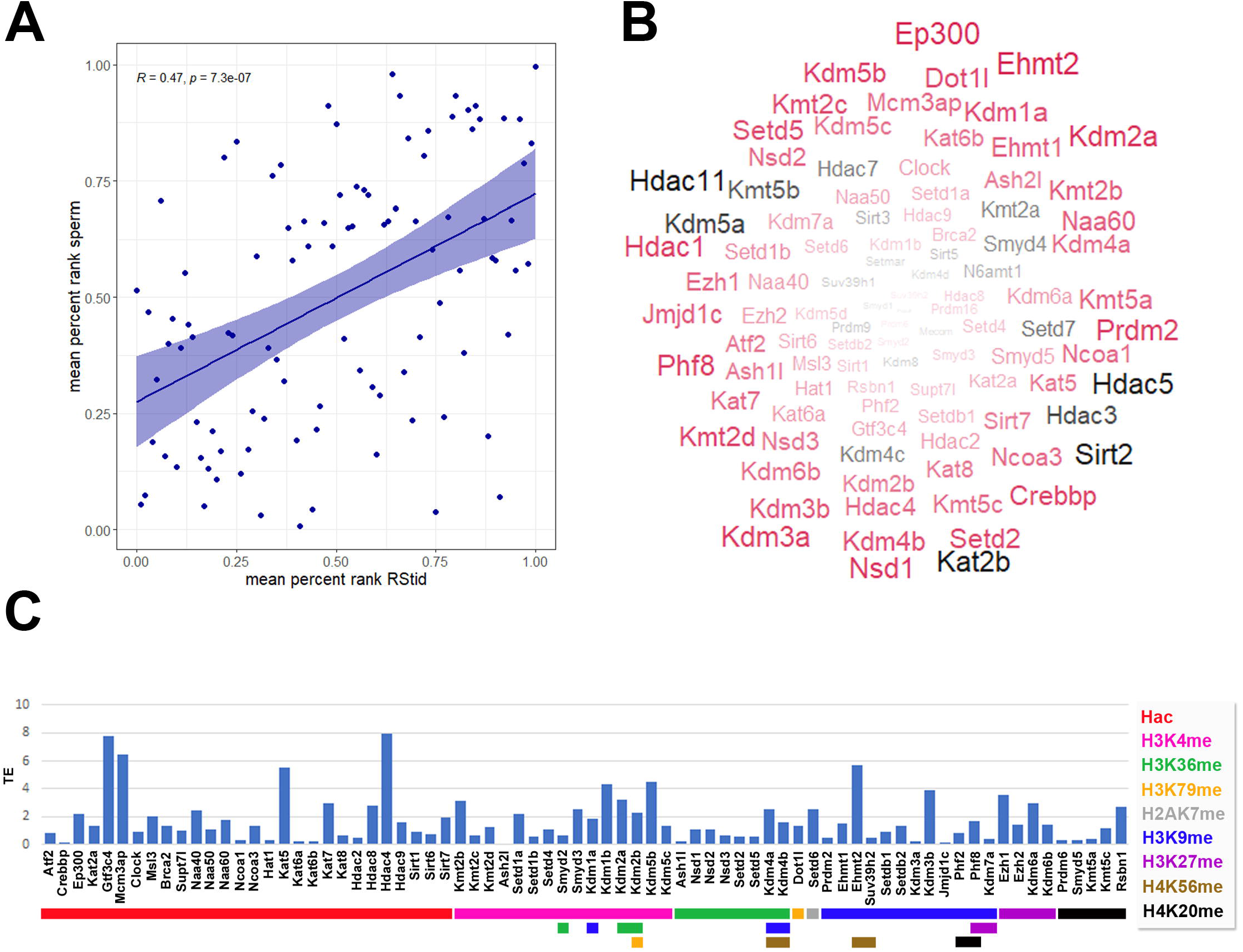
Epigenetic histone K writers and erasers detected at mature sperm RNA cargo and 1-cell embryo translatome. A) Pearson correlation between epigenetic enzymes mean percent rank detected at RStid and sperm. B) Mature sperm word cloud for epigenetic enzymes indicating the enzyme abundance (word size = mean percent rank) and translation efficiency (TE) detected at 1 cell embryo (red: TE > 0, black: TE=0). C) Epigenetic enzymes TE in mouse pre-implantation embryos (TE = RNA RPKM at ribosome / total RNA RPKM, RPKM>0.5)

## DISCUSSION

Accumulating evidence has put on focus the existence of windows of vulnerability during spermatogenesis, at which environmental stressors can induce epigenetic reprogramming and the transmission of developmental, metabolic, and behavioral traits to offspring. To further elucidate the mechanisms driving histone PTMs changes during the spermatogenic process, we analyzed the epigenetic enzyme expression programs across germ cell differentiation to mature sperm. We show that cell transition across spermatogenesis is characterized by the upregulation of specific histone K methylation and acetylation writers and erasers driving the epigenome changes necessary for spermatogonia differentiation, meiosis entry and spermiogenesis. Moreover, most of these enzymes are detected in the mature sperm mRNA pool, and some are also detected in the early zygote translatome, suggesting they can be delivered and participate in the 1-cell embryo chromatin assembly. Our study shows important mechanistic aspects behind transgenerational epigenetics, where epigenetic enzymes not only can respond to environmental stressors and alter the germ cell epigenome but could function as vectors of epigenetic information transmission themselves, by participating in chromatin organization at the maternal-to-zygote transition.

The spermatogonia stage seems to be vulnerable for stress-induced epigenetic reprogramming since they reside outside the blood-testis barrier (González et al., 2020). Furthermore, current data support a dynamic stem cell model in which the fate of Sgund population is context-dependent and plastic, and several epigenetic mechanisms are implicated in the regulation of spermatogonia maintenance and cell fate (Mäkelä et al. 2019). Here, we found that the transition to the SGdiff population is characterized by increased expression of writers of activation marks H3K4me and H3K36me and repression marks H3K9me and H4K20me. At the spermatogonial phase, we found high expression of H3K4me epigenetic enzymes, including *Kmt2a/b/c* writers and *Kdm1a/b* erasers

In this context, at spermatogonial stage occurs the deposition of monovalent and bivalent H3K4me marking at promoters by *Kmt2b* (*Mll2*), preparing germ cells for gene activation at late spermatogenesis and embryonic development (Tomizawa et al., 2018). Accumulating evidence also points to H3K4me2 in the maintenance of transcriptional states during cell development and, its removal by erasers expressed at spermatogonia such as KDM1A (LSD1) is a key step for epigenetic reprogramming and cell fate (Martinez-Gamero et al., 2021). KDM1A was found to mediate spermatogonia commitment and differentiation, as testicular deletion of *Kdm1a* leads to progressive germ cell loss, downregulation of self-renewal factors and abnormal accumulation of meiotic spermatocytes (Lambrot et al., 2015; Myrick et al., 2017). Interestingly, overexpression of *Kdm1a* altered the H3K4me2/3 levels and the RNA profile of sperm and embryos transgenerationally (Siklenka et al., 2015; Lismer et al., 2020). Hence, the spermatogonial phase seems to be especially sensitive to alterations in H3K4me balance linked to epigenetic inheritance. SGdiff showed a specific increase of *Prdm9*, responsible for H3K4me3 and H3K36me3 that mediate the double-strand breaks (DSBs) formation at meiotic hot spots (Sun et al., 2020; Getun et al., 2017). In line with this, it has been shown that meiosis entry of SGdiff is driven by increased levels of H3K36me3 (Godmann et al., 2007; Song et al., 2011; Zuo et al., 2018). Here, we found *Setd2* expression until PD stage, and *Setd5* specifically upregulated at SGund, suggesting their involvement in the specific H3K36me3 landscape of spermatogonia differentiation. The SGund population also expressed H3K4me/H3K36me eraser *Kdm2b*, a subunit of non-canonical PRC1.1 complex that was found to protect the polycomb-silenced promoters against ectopic de novo H3K36me methylation (Blackledge and Klose, 2021). Furthermore, KDM2B was recently found to erase H3K79me2/3 and to induce transcriptional repression via SIRT1 chromatin silencing (Kang et al., 2018).

It is established that SGund typically lacks heterochromatin, which forms as they differentiate to the spermatogonia population committed to meiosis (Chiarini-Garcia et al., 2002). The SGund showed increased expression of H3K9me erasers *Kdm3a/b* and *Jmjd1c* (*Kdm3c*), that were found to counteract the enzymatic activity of EHMT2 (Tachibana et al., 2007), which is highly expressed in the spermatogonia population. Methylation of H3K9 by EHMT2 blocks gene expression of the stem cell factors OCT4 and NANOG, therefore, demethylation of H3K9 by KDM3 family enzymes may be a key step in the maintenance of self-renewal of spermatogonial stem cells (Kuroki et al., 2020; Chioccarelli et al., 2020). In agreement, we found increased *Ehmt1, Suv39h2* and *Setdb2* expression at SGdiff stage, consistent with H3K9me2/3 deposition during spermatogonia differentiation. In particular, EHMT1 was found in a complex that binds E2F- and Myc-Responsive genes to repress the mitotic program (Ogawa et al., 2002). Moreover, SGdiff population showed decreased *Ezh1* and increased *Kdm6b* compared to SGund, suggesting that spermatogonia differentiation involves H3K27me removal. We also observed that the transition to the SGdiff population is characterized by specific HDACs upregulation. In this sense, it was shown that class IIa HDACs enzymatic activity depends on their recruitment into a complex containing class I HDAC3 and retinoid acid (RA) receptor (RAR), suggesting that these HDACs could mediate chromatin silencing induced by RA, essential for the transition of SGund into SGdiff (Parra, 2015). Moreover, histone acetylation writer *Kat2b* expressed in SGund functions as a co-activator for RAR and can interact with NCOA1 to promote transcription (Spencer et al., 1997), which is upregulated from SGdiff to LZ stage. SGund also showed peak expression of *Kat2a*, and its germ-cell-specific knockout results in abnormal chromatin dynamics, leading to increased sperm histone retention and severe reproductive phenotype (Luense et al., 2019). Finally, SGdiff population showed increased *Sirt1*, described as a key regulator of spermatogonia differentiation since SIRT1-deficient mice show a delay of pre-meiotic differentiation, aberrant expression of spermatogenic genes, abnormal spermatozoa with elevated DNA damage, and reduced fertility (Coussens et al., 2008; Bell et al., 2014). Therefore, the action of certain epigenetic enzymes in pre-meiotic stages seems to enable the post-meiotic transcriptome and chromatin remodeling processes.

Initiation of male meiosis is one of the most important events that coincide with spermatocyte differentiation, and meiotic prophase is accompanied by several alterations of epigenetic and gene expression programs for post-meiotic spermiogenesis (Maezawa et al., 2020). Histone acetylation writers increase during prophase I progression, and H3K9/18/23ac and H4K5/8/12/16/91ac marks have been associated with open chromatin and hot spot cores (Getun et al., 2017; Chioccarelli et al., 2020). Accordingly, MYST enzymes were previously found strongly related to meiosis control, increasing their expression from early pachytene through diplotene stages (Getun et al., 2017). Interestingly, p300/CBP family and KAT2B were found to regulate histone eviction by acetylation of transition nuclear protein (TNP) 2, affecting DNA condensation properties and interaction with histone chaperones (Pradeepa et al., 2009). Among other acetylation writers upregulated during prophase I, MSL3 binds to H3K36me3 marked sites and deposits H4K16ac, controlling meiosis entry and STRA8 (stimulated by retinoic acid 8) functions (McCarthy et al., 2022). On the other hand, we detected several SIRT enzymes throughout prophase I. Deacetylation of H3K9 by SIRT6 modulates telomeric chromatin function (Michishita et al., 2008) while SIRT7 is highly selective toward H3K18ac, and might play an upstream role in DNA repair and telomere maintenance (Wu et al., 2018). We also detected peak expression of *Hdac1* and *Kdm1a* (*Lsd1*) at the PD stage, that were recently shown to interact with BEND2 and participate in DSB repair, synapsis and transcriptional repression (Ma et al., 2022). HDAC1 was also found to form a complex with DNA methyltransferase DNMT3L to regulate X chromosome compaction (Deplus et al., 2002; Zamudio et al., 2011).

During prophase I, recombination hotspots are mainly marked by H3K4me3 catalyzed by KMT2 family and PRDM9 methyltransferases (Sollier et al., 2004; Buard et al., 2009). Moreover, the levels of H3K4me3 and H3K36me3 are highly correlated at hotspots, but mutually exclusive elsewhere, and PRDM9 is capable of placing both marks on the same nucleosomes in vivo (Powers et al., 2016). It was shown that *Prdm9* knockout changes the distribution of DSBs across the genome inducing defective synapses and male infertility (Paigen and Petkov, 2018; Bhattacharyya et al., 2019). We also detected peak expression of H3K79me writer *Dot1L* at PD stage, consistent with previous reports showing increased levels of DOT1L, and H3K79me2/3 from pachytene onwards (Ontoso et al., 2014). The heterochromatic centromeric regions and the sex body are enriched of H3K79me3, while H3K79me2 is present all over the chromatin, but is largely excluded from the sex body despite the accumulation of DOT1L (Ontoso et al., 2014). Repressive histone methylation marks H3K27me3, H3K9me3 and H4K20me3 are present in chromatin regions with reduced recombination or involved in heterochromatin formation. *In vitro* and *in vivo* models disrupting the action of H3K9me writers *Suv39h1, Setdb1/2 and Ehmt2* detected during prophase I, showed an abnormal distribution of H3K9me3 at pericentromeric heterochromatin, compromised meiotic silencing of unsynapsed chromatin (MSUC), misregulation of meiotic and somatic genes, anomalous synapsis and misssegregation of chromosomes and apoptosis at pachytene stage (Peters et al., 2001;, Tachibana et al., 2007; Takada et al., 2011). Interestingly, low protein diet in the father was linked to altered levels of H3K9me2 through EHMT2, that changed tRNAs in sperm and transmitted metabolic phenotypes to the offspring (Yoshida et al., 2020). Also, SETDB1 haploinsufficiency in mice was shown to induce changes in DNA methylation in transposable elements, and to influence coat color in the offspring, further linking epigenetic inheritance with altered epigenetic enzymes levels during spermatogenesis (Daxinger et al., 2016). A recent work by Barral et al., 2022 reported dual regions in mouse embryonic stem cells that rely on the SETDB1 and NSD proteins to generate H3K9me3 and H3K36me3, respectively. They found that SETDB1 removal induces loss of both marks in dual regions, gains signatures of active enhancers, and comes into contact with upregulated genes, providing a mechanistic insight by which genes are controlled by heterochromatin (Barral et al., 2022). Moreover, NSD1-mediated H3K36me2 was found to prevent H3K27me3 deposition by PRC2, modulating PRC2-mediated H3K27me domains demarcation (Blackledge and Klose, 2021). In line with this, we detected the expression of H3K27me writers *Ezh1/2* at PreL and LZ stages that were found involved in mammalian X chromosome inactivation (Schuettengruber et al., 2007). Furthermore, *Kdm6a* is part of the COMPASS complex with *Kmt2b/c/d* and *Ashl2* that activate genes in response to RA by H3K4me deposition and H3K27me removal (Lavery et al., 2020) and moreover, alterations in *Kdm6a* expression induce defects that persisted transgenerationally (Siklenka et al., 2015, Lesch et al., 2019).

Additionally, the loss of H3K27me3 from bivalent signature in the promoter of endonuclease SPO11 induces chromatin activation that favors meiotic entry (Hammoud et al., 2014). We also detected high expression of H2AZK7me1 writer *Setd6* at PreL stage, which in ESC was found with H3K27me3 close to differentiation marker genes and removed upon RA signal (Binda et al., 2013), suggesting that SETD6 may have a role at early male meiosis. Finally, we observed expression of the H4K12me writer *N6amt1* at LZ stage, a recently described epigenetic mark found at promoters of genes encoding cell cycle regulators (Metzger et al., 2019).

Chromatin organization dramatically changes during mid-to late-spermiogenesis, due to histone eviction and replacing by PRMs to facilitate the condensation and packaging of the paternal genome. In mice, 1%-10% of histones are retained in sperm chromatin and form a heterogeneous mixture of nucleo-histones and nucleo-protamines (Rajender et al., 2011; Luense et al., 2016). At the spermiogenesis phase, histone acetylation is essential in destabilization and remodeling of nucleosomes. Here, the RStid population showed high transcriptional levels of H4K16ac writers *Msl3* and *Kat8*, a histone PTM essential for histone eviction which does not occur in the absence of this specific mark (Lu et al., 2010). Also, *p300/CBP* and *Kat2b* keep their expression from spermatocyte stage, due to their crucial role to regulate histone eviction (Pradeepa et al., 2009). Moreover, KAT2B was found in spermatids and involved in H3K9ac, an epigenetic mark detected at unmethylated active genes / enhancers (Hammoud et al., 2014) that could influence gene expression directly after fertilization (Steilmann et al., 2011). Additionally, *Hat1* expression was found related to the incorporation of H4K5/12ac and H3.3 variant at DSBs sites and the promotion of DNA repair (Yang et al., 2013). H4K5/8/12 marks have been also observed just before eviction of histones during spermiogenesis (Meistrich et al, 1992; Hecht et al., 2009) and moreover, H4K8/12ac was detected prior to full decondensation of the sperm nucleus, suggesting that these marks are transmitted to the zygote (van der Heijden et al., 2006).

Multiple histone methylation have been also identified in spermatids, pointing to a balance of “opened” and “closed” chromatin regions during the histone-to-protamine transition (Wang et al., 2019). Here, we detected upregulated expression of H3K79me writer *Dotl1* and H3K9me writer *Ehtm2* at RStid stage. It was found that DOTL1 is enriched in post-meiotic stages of mouse germ cells and precedes the histone-to-protamine transition (Dottermusch-Heidel et al., 2014). During spermiogenesis, there is a reactivation of transcription from MSUC and MSCI sites that is enabled by the deposition of histone crotonylation (Kcr). It was proposed that CDYL, an HKcr eraser, prevents post-meiotic chromatin reactivation by binding to H3K9me3 and H3K27me2/3 marked sites and facilitating H3K9me2 deposition by EHMT2 (Mulligan et al., 2008). Moreover, there is specific H3K27me deposition to establish bivalency at developmental genes (Maezawa et al., 2018), and proteomic studies found substantial H3K27me3/H3K36me2 double marking in RStid and sperm (Luense et al., 2016).

It is established that mature sperm transport a cargo of miRNAs, tsRNAs, lncRNAs, circRNAs, and protein-coding mRNAs, that carry an epigenetic blueprint involved in early embryo development (Guo et al., 2017). Some of these RNAs remain from the last stages of elongated spermatids, and others are acquired along the passage through the epididymis, as sperm absorb epididymosomes released by somatic cells (Trigg et al., 2019). Here, we found that the mature sperm transport a signature of epigenetic enzymes mRNAs similar to the one detected at spermiogenesis phase, and most of these enzymes show positive translation at 1 cell embryo. After fertilization, the zygote genome is transcriptionally silent, and cellular processes are carried on with inherited RNAs and proteins until the onset of zygotic genome activation (ZGA) around the 2-4 cells stage (Schulz and Harrison, 2019). Also, at 1 cell stage there is significant removal of some maternal RNAs and specific translation of others, including RNAs specifically delivered by the sperm like the egg-activating factor PLC-zeta (Sone et al., 2005; Yao et al., 2010). The parental pronuclei have asymmetric reprogramming capacities and the reprogramming factors reside predominantly in the male pronucleus. Here, we detected high sperm values and embryo translation of several enzymes involved in histone acetylation and methylation, including the H4K20me2/3 writer *Kmt5c*, the H3K9me2 writers *Ehmt1/2*, and the H3K9me3 writers *Suv39h2* and *Setdb1/2*, with established roles in chromatin structure organization. In this context, it was found that EHMT2 activates soon after fertilization and deposit H3K9me2 patterns in the paternal genome (Ma et al., 2015). Also, previous work suggested that KMT5C deposits H4K20me3 to allow the timely and coordinated progression of replication after fertilization (Eid et al., 2016).

## CONCLUSIONS

In summary, the analysis performed here shows important windows during spermatogenesis, where interference with epigenetic enzymes gene expression may have phenotypic consequences in the offspring, even when they are not inherited. Moreover, we show that epigenetic enzymes mRNA could become functional in pre-cleavage zygotes and contribute to early chromatin organization with deep implications in future embryo development. The epigenetic enzyme’s paternal contribution could be another mechanism for epigenetic inheritance that deserves further consideration.

## Data availability

The datasets analyzed during the current study, enzymes metadata table and full R scripts are available at https://github.com/Gonzalez-Lab/Gonzalez-2022-germ-cells.

## Competing interests

The authors declare no competing interests.

## Funding

This work was supported by Agencia Nacional de Promoción Científica y Tecnológica (PICT 2019-00171, BG) and Fundación Científica Felipe Fiorellino (AV and CG).

## Authors’ contributions

B.G and C.R.G conceived the project, analyzed the data and wrote the manuscript. G.B, M.S.F and C.P.L contributed to data mining and analysis. A.D.V contributed to the critical reading and editing of the manuscript. All authors read and approved the submitted version.

## Notes

https://github.com/Gonzalez-Lab/Gonzalez-2022-germ-cells

### Competing Interest Statement

The authors have declared no competing interest.

### Summary of Updates

We found an error on the previous version, a figure was repeated and the corresponding figure was missing.

