## Supplemental info for "Transcriptome profiling of histone writers/erasers enzymes across spermatogenesis, mature sperm and pre-cleavage embryo: Implications in paternal epigenome transitions and inheritance mechanisms"

Supplemental information

Dataset validation

To validate the GSE dataset, we evaluated consistent markers expression across cell types. SGund showed peak expression of stem cell markers *Thy1*, *Nanos2*, *Nanos3*, *Gfra1*, *Zbtb16* and *Uchl1* and SGdiff the committed spermatogonia marker *Kit*. As it is established, the transition from SGund to SGdiff coincides with the gain of the c-kit receptor in the adult testis. Kit continues to be expressed until meiosis and plays essential roles in the survival of SGdiff. The PreL stage showed shared expression with SGdiff gene *Kit* and LZ stage genes *Dazl* and *Top2a*. LZ stage showed differentiation markers *Stra8* and *Sycp1/2/3*, whereas PD stage showed meiosis markers *Spo11*, *Piwil* and *Pttg1*. The PD stage shared gene expression with RStid of *Ddx4*, *Dkkl1*, *Insl6*, *Acrbp* and *Acr*, and RStid stage also showed peak expression of classical spermiogenesis markers *Tnp1/2* and *Prm1/2*, and *Izumo1*, *Spag6*, *Acrv1* and *Pgk2*. To check the germ cells purity, we analyzed marker genes of other cell populations in the testis including Leydig (*Cyp11a1*, *Cyp17a1*, *Hsd3b1* and *Star*), Sertoli (*Sox9*, *Drd4* and *Rhox8*), endothelial (*Vwf* and *Tie1*), myoid (*Acta2* and *Myh11*) and machophage (*Adgre1*) cells, which were detected in very low levels (supplemental Table 1).

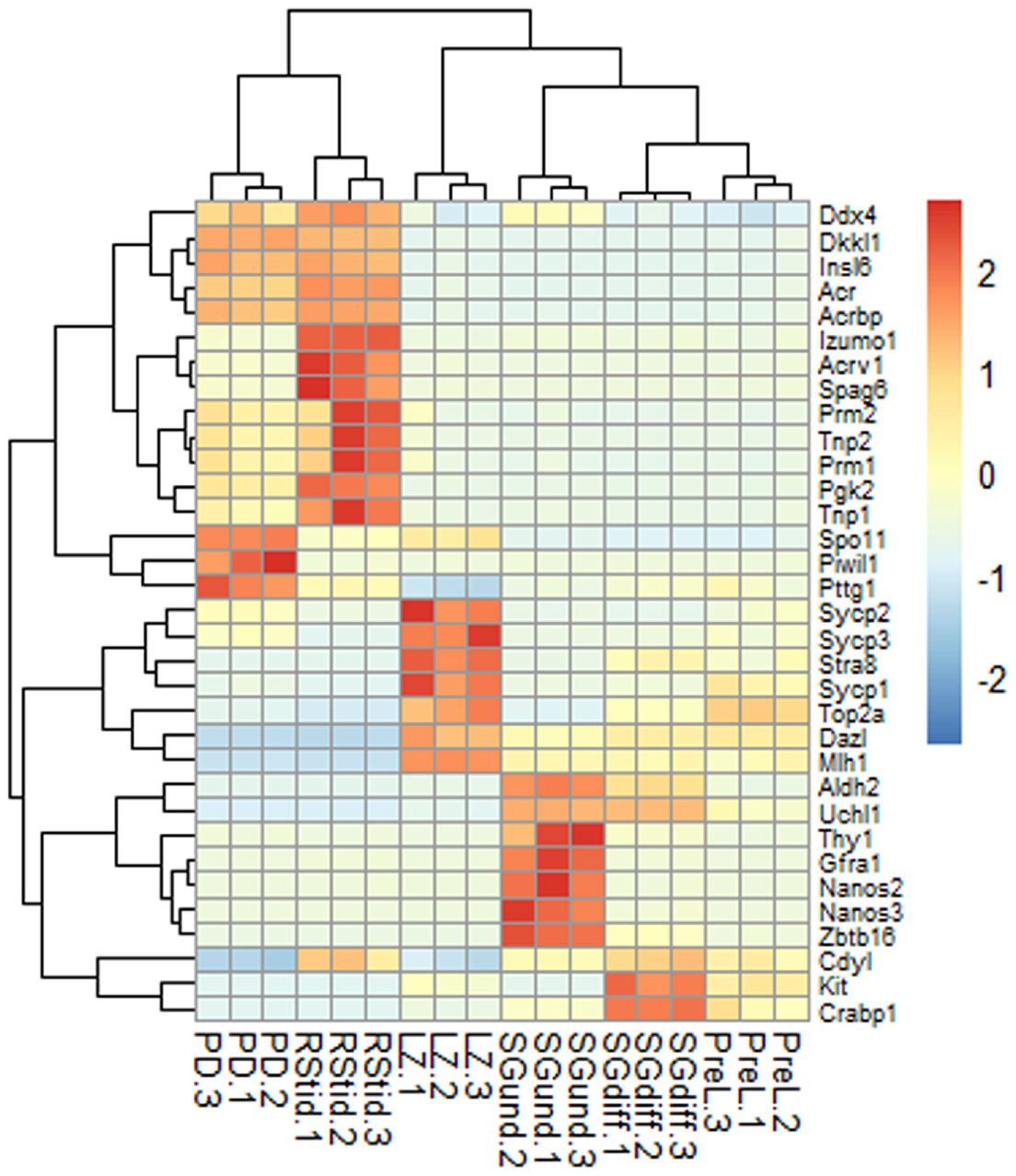
